# Metabolic Cage Analysis of Surgically Catheterized C57Bl/6J Mice (*Mus musculus*) Treated with Carprofen and Sustained-release Buprenorphine

**DOI:** 10.1101/2025.05.21.655398

**Authors:** Jui Rose Tu, Marissa Saenz, Elizabeth Bloom-Saldana, Patrick T. Fueger

## Abstract

Federal regulations require that appropriate analgesia be provided for laboratory animals for pain control. Carprofen and buprenorphine are two common analgesics used for laboratory mice (*Mus musculus*). However, given the potential gastrointestinal side effects that these analgesics have in various species, the impact of these analgesics on mice used in metabolic studies could be concerning. To investigate the impact of carprofen and sustained-release buprenorphine on food consumption, activity level, and whole-body metabolism, we administered carprofen alone or in combination with sustained-release buprenorphine to mice that underwent jugular vein and carotid artery catheterization, or a sham surgery. The mice were individually housed in instrumented metabolic cages to continuously quantify food consumption, activity levels, and energy expenditure by indirect calorimetry. We hypothesized that catheterized mice receiving both carprofen and sustained-release buprenorphine would have decreased food consumption and increased activity level compared to mice that received sham surgery and carprofen, and catheterized mice treated with carprofen only would have similar food consumption and activity level as sham mice that received carprofen. Our results demonstrate that during the initial 12h after surgery, catheterized mice that received both carprofen and sustained-release buprenorphine were more active than sham mice that received carprofen, and were more active and consumed more food than catheterized mice that received carprofen only. Our study demonstrated how analgesia regimen can affect metabolic parameters. Therefore, researchers should carefully consider the effects that analgesic drugs can have on mice when designing metabolic or behavioral experiments.

## Introduction

The *US. Government Principles for the Utilization and Care of Vertebrate Animals Used in Testing, Research, and Training* states that “…unless the contrary is established, investigators should consider that procedures that cause pain or distress in human beings may cause pain or distress in other animals.”^3^ The *Guide for the Care and Use of Laboratory Animals* further emphasizes that managing pain and distress in laboratory animals is an integral component of veterinary medical care.^4^ Opioids and nonsteroidal anti-inflammatory drugs (NSAIDs) are frequently used in laboratory animal species, and they are often used together to achieve multi-modal pain control.^5^ In the laboratory setting, the design of pain control regimen should take into consideration of the species, procedures, and research needs.^4,6^ Whereas NSAIDs have little to no reported behavioral effects on laboratory rats and mice, studies suggest that opioids may have stimulatory effects on mice,^7,8^ and cause pica in rats.^9^

The opioid buprenorphine has long been used in human and veterinary medicine. First synthesized in 1966, buprenorphine was initially registered as an analgesic in the UK, and then worldwide in the 1980s.^1^ In 2002, buprenorphine-based products were approved by the FDA as treatments for opioid use disorder.^2^ Since then, additional formulations of buprenorphine products have become available, including products for minor species such as rats and mice. Buprenorphine’s long history of medical use had led to the development of substantial clinical knowledge on its use and actions. Yet in veterinary medicine, much remains to be learned regarding buprenorphine’s effects in different species.

Buprenorphine is thought to affect activity levels of mice, as previous studies have reported that opioid administration to mice is associated with hyperactivity.^10,11^ In terms of food consumption, opioids can bind to the opioid receptors in the gastrointestinal (GI) tract and cause clinical signs such as indigestion, nausea, and vomiting in humans.^12^ Comprehensive evaluation of the effect of time-release buprenorphine formulations on the metabolic parameters of mice is lacking, although such information is essential to know for both behavioral and metabolic studies. To address this critical gap in knowledge, we designed the following experiment: males and female C57BL/6J mice received either jugular vein and carotid artery catheterization surgery, or sham surgery. Sham mice received carprofen for analgesia (Sham+C), and catheterized mice received either carprofen only for pain management (Sx+C), or both carprofen and sustained-release buprenorphine (Sx+C+SRB). The metabolic phenotype of the mice was recorded and evaluated before and after the surgery with an instrumented metabolic caging platform capable of monitoring food consumption, activity, and energetics by indirect calorimetry. We hypothesized that mice the in Sx+C group would have similar food consumption and activity level as mice in the Sham+C group, and mice in the Sx+C+SRB group would have decreased food consumption and increased activity level compared to the Sham+C mice (**Fig. 1**).

**Figure 1.**
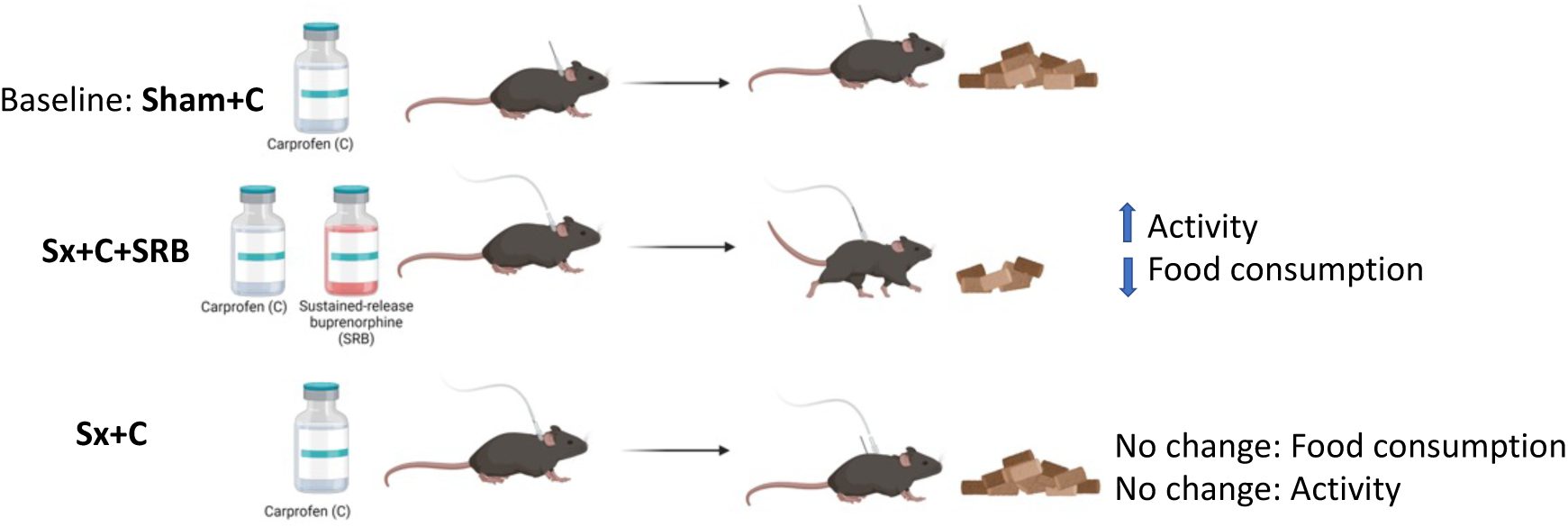
Study design and hypothesis. Mice were subjected to baseline measurements and then were assigned to one of three treatment group: sham surgery plus carprofen (Sham+C), catheterization surgery plus carprofen and buprenorphine (Sx+C+SRB), and catheterization surgery plus carprofen alone (Sx+C). Post-surgical data were collected and the a priori hypotheses are indicated.

## Material and methods

Many of the material and methods have been described in detail in a related publication by Saenz *et al*.^13^ and are summarized as follows: All animal procedures were approved by the IACUC at the City of Hope National Medical Center (Duarte, CA), and performed on site in an AAALAC-accredited facility. Six-week-old male (*n* =18) and female (*n* = 29) C57BL/6J mice were purchased from The Jackson Laboratory (Bar Harbor, ME). The mice were acclimated to the vivarium until 13-15-week of age, and then underwent catheterization surgery. During the acclimation period, the mice were group-housed in Optimice individually ventilated cages (Animal Care systems, Centennial, CO) with corncob bedding (Bed-o’-Cobs, 1/8-in, The Andersons, Maumee, OH), cotton nestlet, and PVC tube. Mice had *ad libitum* access to rodent chow (no. 5053, LabDiet, St. Louis, MO) and reverse-osmosis-purified water. Three days before the surgery, mice were handled daily for about 3 min, and provided with diet gel (Dietgel Recovery, Clear H2O, Westbrook, ME). Diet gel was also provided to all mice for 3 d after surgery. The light:dark cycle was 12:12h, and the light was provided by recessed fluorescent lighting fixtures emitting 35-44 lux at the level of the cage. The health status of the mice was SPF, free of the following pathogens: mouse rotavirus, Sendai virus, pneumonia virus of mice, mouse hepatitis virus, minute virus of mice, mouse parvovirus, Theiler murine encephalomyelitis virus, mouse reovirus type 3, mouse norovirus, lymphocytic choriomeningitis virus, mouse thymic virus, mouse adenovirus types 1 and 2, mouse cytomegalovirus, polyoma virus, K virus, ectromelia virus, Hantavirus, LDH-elevating virus, *Pneumocystis* spp*., Bordetella* spp., *Corynebacterium bovis, Corynebacterium kutscheri, Campylobacter genus, Helicobacter* spp*., Klebsiella* spp*., Streptobacillus moniliformis, Salmonella* spp., *Staphylococcus aureus, Streptococcus pneumonia,* Beta *Streptococcus* spp., CAR *bacillus, Encephalitozoon cuniculi, and Mycoplasma pulmonis, Helicobacter* spp*., and Clostridium piliforme,* endo- and ectoparasites.

### Experimental design

Mice were randomly divided into 3 treatment groups, and males (M) and females (F) were distributed as evenly as possible. Some mice failed to thrive after the catheterization surgery. The final group sizes were as follows: Sham+C, *n* = 16 (10F, 6M); Sx+C, *n* = 16 (10F, 6M); Sx+C+SRB, *n* = 15 (9F, 6M). The Sham+C mice received a sham surgery, then received carprofen (5 mg/kg, subcutaneously, once daily for 3 d) for analgesia; the Sx+C mice received catheterization surgery of a carotid artery and jugular vein, then received carprofen (5 mg/kg, subcutaneously, once daily for 3 d) for analgesia, the Sx+C+SRB mice received the same catheterization surgery, and then received both carprofen and one subcutaneous injection of SRB (1 mg/kg Buprenorphine-ER, Zoopharm, Windsor, CO) for analgesia. To ensure accuracy, analgesia was delivered with Hamilton syringes fitted with 23-gauge needles.

Metabolic parameters were acquired with instrumented metabolic cages (Promethion Core Metabolic System, Sable Systems, Las Vegas, NV). Two days prior to catheterization surgery, mice were housed individually in the metabolic caging platform for baseline recordings. After the surgery, mice were housed in the same metabolic cages for 3 d for subsequent data acquisition. Pain was assessed by performing cage side observation with a modified rubric (Table-1) from Adamson *et al.*^14^ Pain scoring was performed before surgery and then at 0.25, 1, 2, and 3 d post-surgery.

### Surgical procedure

The surgical procedure for carotid artery and jugular vein catheterization has been previously described in detail.^15^ Mice were anesthetized with isoflurane, then the ventral cervical and dorsal intrascapular regions were clipped and scrubbed to prepare for aseptic procedures. A longitudinal incision about 5-mm was made at the ventral cervical region, slightly left to the trachea. The left common carotid artery was isolated and catheterized with a silastic-polyethylene 10 catheter (Fisher scientific, Waltham, MA). The catheter was prefilled with 100 U of heparin-saline solution. Next, another longitudinal incision was made over the right jugular vein, and then the vein was isolated and catheterized in the same manner. The free ends of the catheters were tunneled subcutaneously toward the dorsal interscapular region, excited via a 5-mm incision, and connected to a “Mouse Antenna for Sampling Access” (MASA™; made inhouse) device. The device serves as the port by which the catheters can be accessed for blood sampling, and when not in use, a cap covers the port. The device was secured on the mouse with suture. For mice that received catheterization surgery (Sx+C, and Sx+C+SRB groups), the above-mentioned procedures were performed. For mice that received sham surgery (Sham+C group), the carotid artery and jugular vein were isolated but not catheterized, and about 10 min later the incisions were closed, and a MASA device was implanted. Based on appearance, a blinded observer was unable to distinguish between treatment groups.

### Data collection and analysis

Raw metabolic cage data were collected by the Promethion and exported as Excel files, which were then analyzed using CalR, a web-based analytical tool for metabolic cage data. Statistical analysis was performed using Graphpad Prism (GraphPad Software, San Diego, CA). Data are presented as means ± SEM.

## Results

### Body weight, food consumption, energy expenditure, and activity level pre-surgery

Pre-surgery, mice were housed individually in the metabolic cages for baseline data acquisition. As expected, between the groups there were no significant differences in body weight, food consumption, energy expenditure, and activity level (as shown in both locomotor activity and distance traveled in cage) (Fig 2a-i). The rise and fall of food consumption, activity level, and energy expenditure closely correlated with the light/dark cycle, where the mice consumed more food and were more active during the dark hours, and the opposite during the light hours.

**Figure 2.**
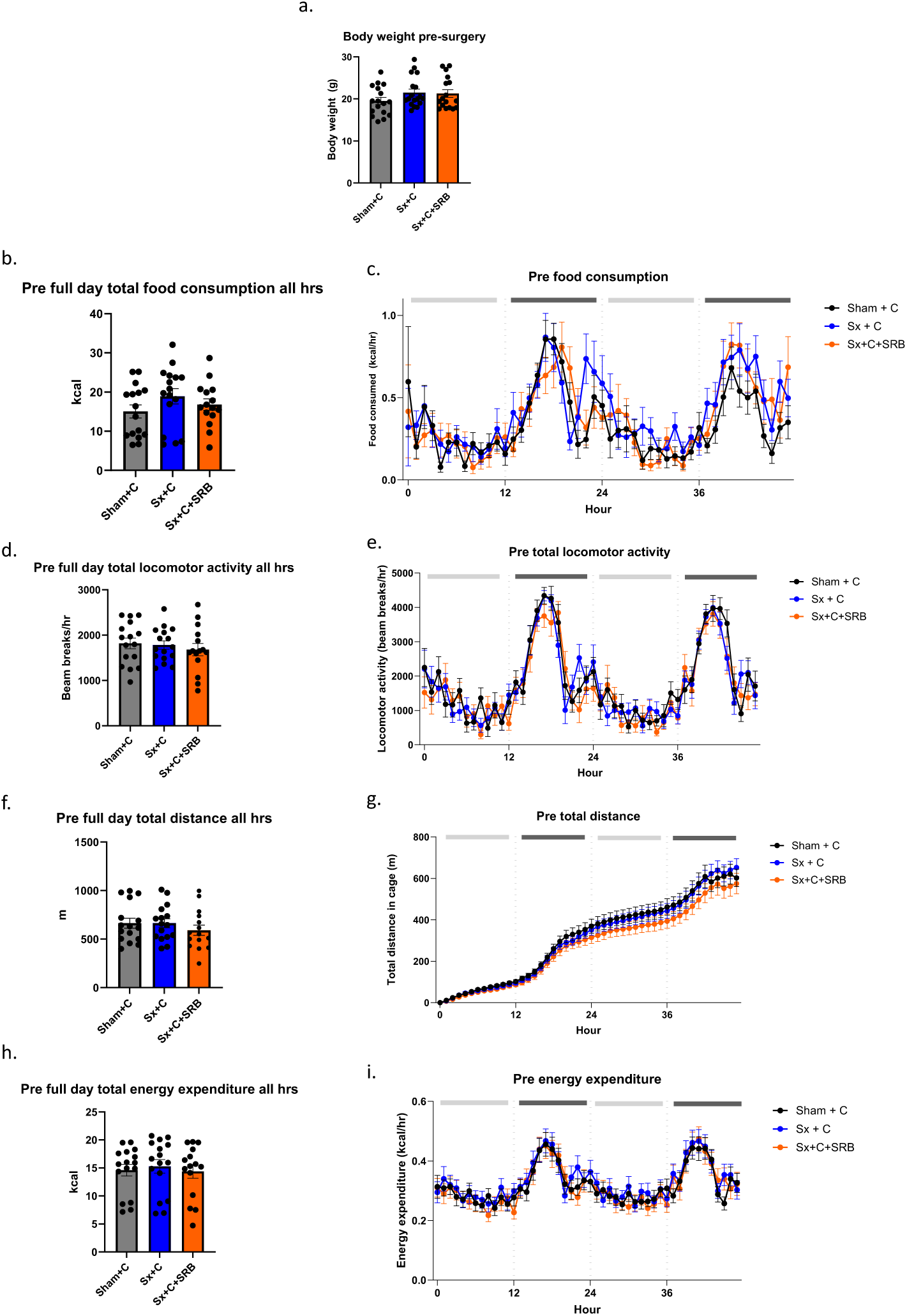
Pre-surgery body weight, food consumption, activity level, and energy expenditure. Prior to any treatment, body weight was measured (A), and mice were placed in instrumented metabolic cages individually to measure food consumption (B,C; in Fig 2C, the light grey bars represent light periods, and the dark grey bars represent dark periods), locomotor activity (D,E), total distance traveled (F,G), and energy expenditure (H,I). Cumulative (B,D,F,H) and continuous (C,E,G,I) data are reported. Data are reported as mean ± SEM; no differences were detected between groups.

### Body weight and food consumption post-surgery

Post-surgery, the mice were returned to their individual metabolic cages for 72 h for data acquisition. The body weights were similar between the 3 groups (Fig. 3a). Contrary to our hypothesis, treatment of SRB did not decrease food consumption. During the 72 h post-surgery, cumulative food consumption was similar between the groups (Fig. 3b), and the rise and fall of food consumption mostly remained correlated with the light/dark cycle (Fig. 3c). However, upon initial examination of the data, during the initial hours post-surgery, the Sx+C group appeared to consume less food (Fig. 3b); therefore we subsequently analyzed the initial 12 h following surgery. With this more focused analysis, the Sx+C+SRB group consumed significantly more food than the SX+C group (Fig. 3d). Next, we lengthened the analysis time period to 24 h to capture one entire light/dark cycle. The results indicated that there was no longer a significant difference in total food consumption between the groups (Fig. 3e). We further analyzed the difference (delta) between post-surgery and pre-surgery food consumption within each group, both during the first 12 and 24 h. The results indicated that Sx+C+SRB was the only group that had a positive delta value during the first 12 h, and the value was significantly higher than that of Sx+C (Fig. 3f). For the 24-h analysis, both Sham+C and Sx+C+SRB had delta values that were significantly higher than Sx+C (Fig. 3g).

**Figure 3.**
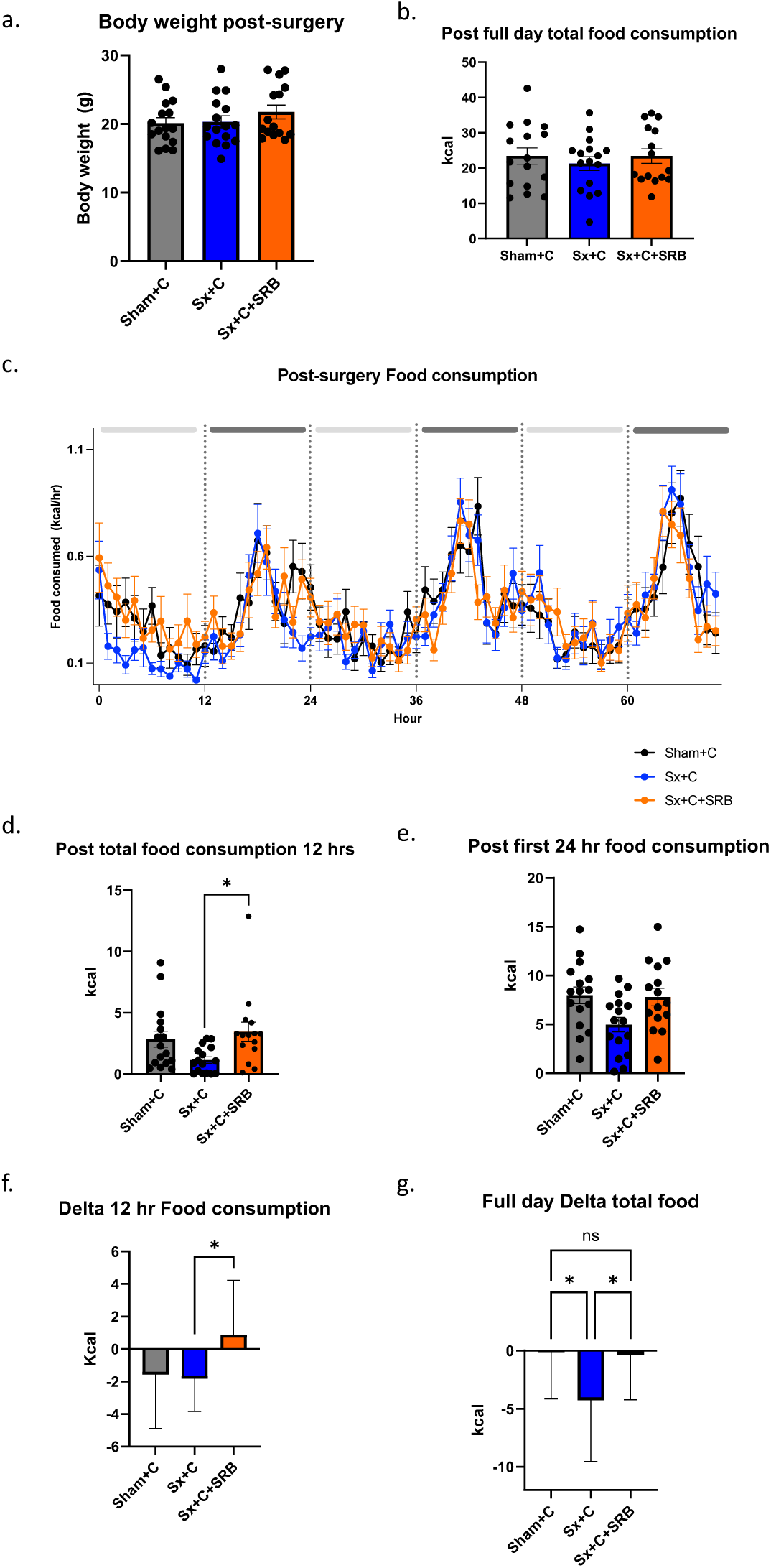
Post-surgery body weight and food consumption. Following surgery and administration of analgesics, mice were placed in metabolic cages for 72 h. Body weight was measured upon removal from the cages (A), and total (B) and continuous (C) food consumption were quantified (B,C); cumulative data over the first 12 (D) and 24 (E) hours are also reported, as are the change in food consumption relative to baseline data (F,G). Data are reported as mean ± SEM; * *p* < 0.05.

The data presented here indicate that SRB treatment did not decrease food consumption. Mice in Sx+C and Sx+C+SRB groups received the same surgical procedure, so it is possible that the lower food consumption observed in Sx+C mice was a result of carprofen alone not providing adequate analgesia during the first 12 h post-surgery.

### Activity level post-surgery

Activity level was captured by the metabolic cages as locomotor activity (beam breaks/hr) and total distance traveled in cage (meters, or m). The cumulative locomotor activity for the entire 72 h post-surgery was similar between the groups (Fig. 4a). The rise and fall of locomotor activity largely followed the light/dark cycle (Fig. 4b), except for the first approximately 12 h post-surgery. Therefore, we analyzed the first 12 and 24 h more closely. This targeted analysis revealed that the Sx+C+SRB group was significantly more active than both the Sham+C and Sx+C groups during the first 12 h (Fig. 4c). However, by 24 h post-surgery, differences were no longer observed amongst the groups (Fig. 4d).

**Figure 4.**
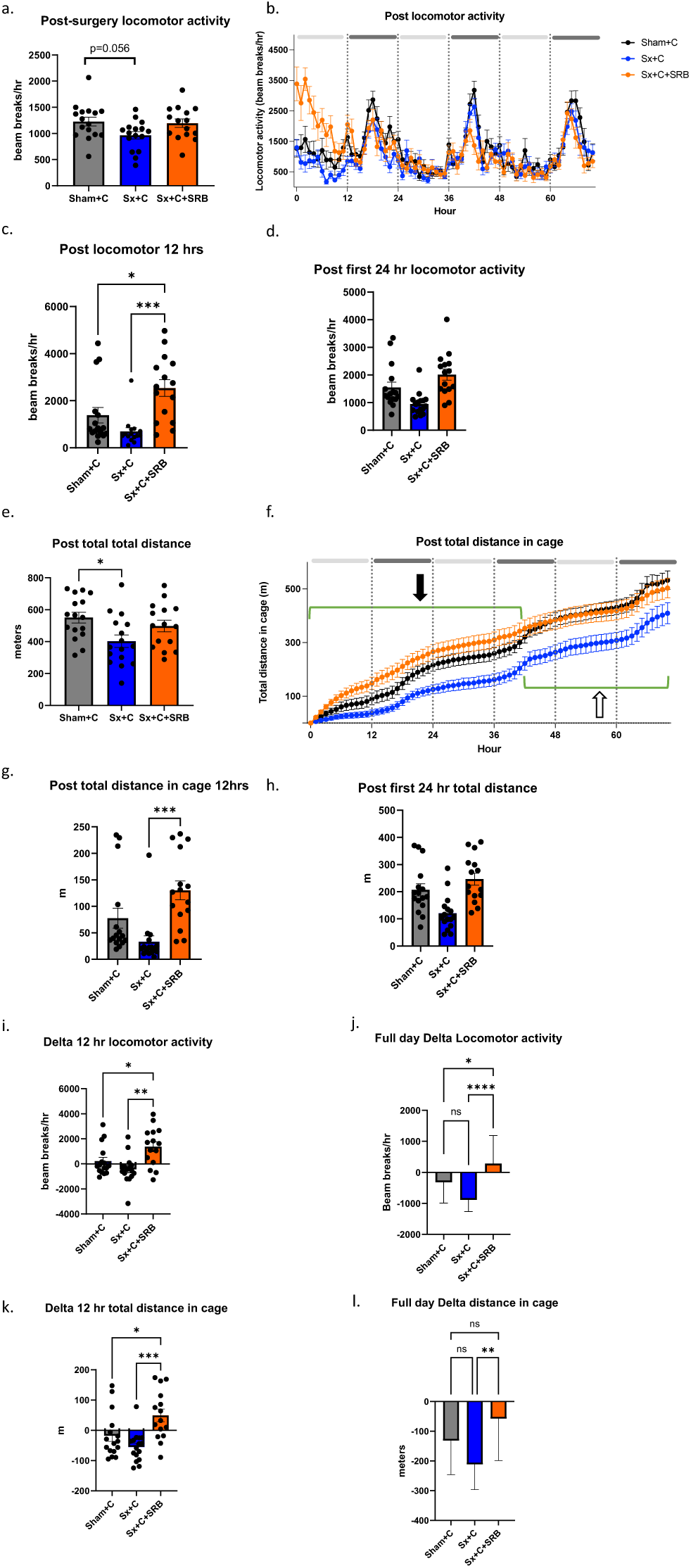
Post-surgery activity level: locomotor activity and total distance traveled in cage. Following surgery and administration of analgesics, mice were placed in metabolic cages for 72 h. Instrumented metabolic caging was used to quantify total locomotor activity (A-D), and distance traveled in the cage over 72 h (E-H); locomotor activity (I,J) and distance traveled in the cage (K,L) relative to baseline data were calculated. Data are reported as mean ± SEM; * *p* < 0.05.

For total distance traveled in cage, the cumulative distance traveled in cage for the entire post-surgical period was significantly lower in the Sx+C compared to Sham+C group (Fig. 4f). The data also indicated that during approximately the first 37 h (Fig. 4f, bracket indicated by the solid arrow), the Sx+C+SRB group had the highest total distance traveled in cage, and then Sham+C started to catch up after that (Fig. 4f, bracket indicated by the open arrow). Time periods of the first 12 and 24 h post-surgery were also analyzed. The results showed that Sx+C+SRB traveled significantly more distance in cage than Sx+C during the first 12 h (Fig. 4g). Then, by 24 h post-surgery, significant differences were no longer observed amongst the groups (Fig. 4h).

Calculation of delta between post- and pre-surgery revealed similar findings in both 12- and 24-h time periods. Sx+C+SRB mice were significantly more active than Sx+C and Sham+C mice (Fig. 4i-k), except for distance traveled in cage during the 24 h period: Sx+C+SRB mice only had significantly higher delta than Sx+C (Fig. 4l). In summary, SRB treatment transiently increased activity level in mice in the initial 12 h post-surgery.

### Energy expenditure

Cumulative energy expenditure level was similar between the groups after the surgery (Fig. 5a), and all 3 groups exhibited higher expenditure during the dark hours (Fig. 5b). Like activity level, Sx+C+SRB appeared to exhibit higher energy expenditure than the other 2 groups during the initial hours post-surgery, therefore we analyzed the first 12 and 24 h further. Importantly, the results showed that there were no significant differences between the groups in terms of cumulative energy expenditure at either timepoint (Fig. 5c-d), thereby indicating that the animals would not require supplemental nutrition to account for an increase in energy usage.

**Figure 5.**
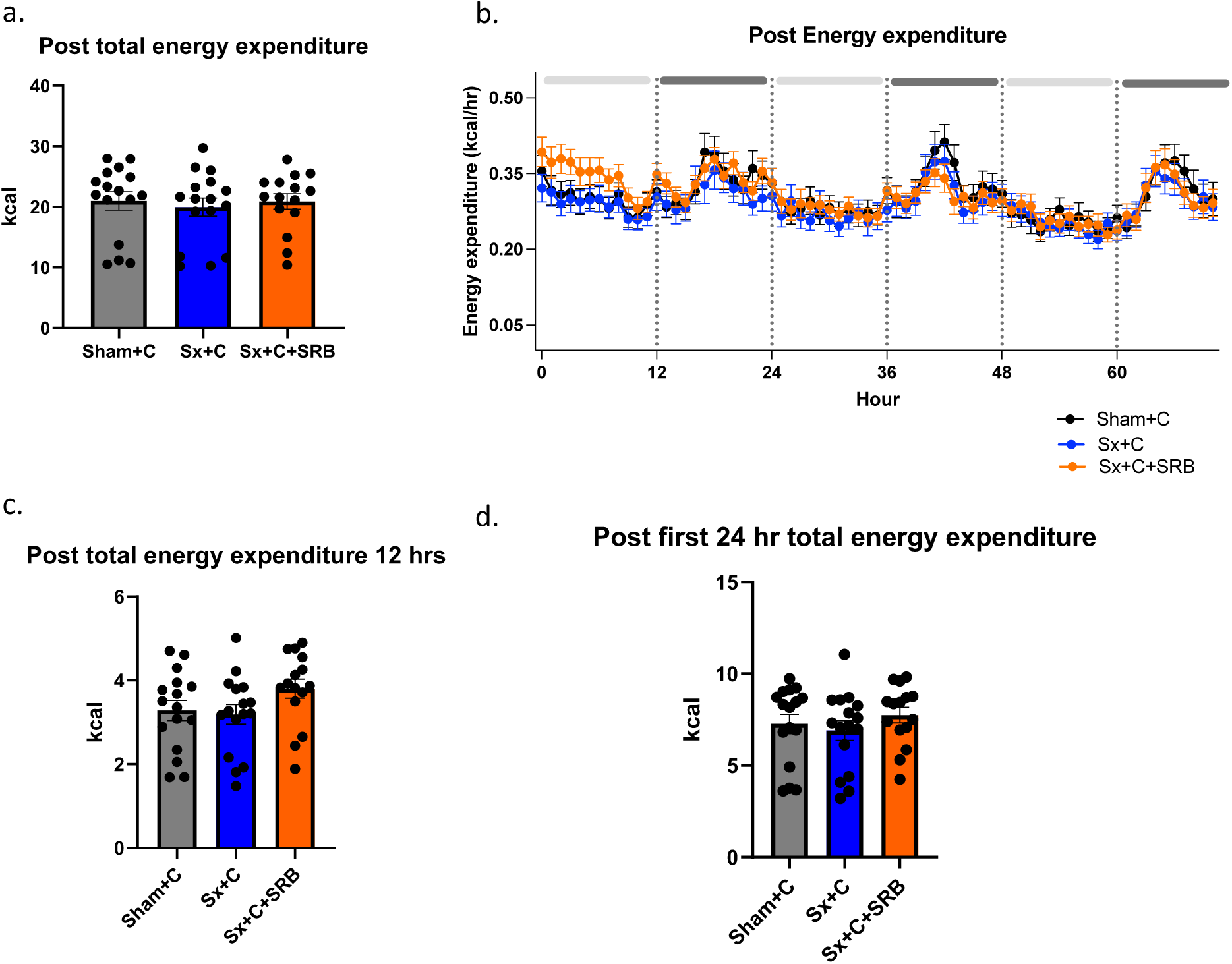
Post-surgery energy expenditure. Following surgery and administration of analgesics, mice were placed in metabolic cages for 72 h. Instrumented metabolic caging was used to quantify total (A) and continuous (B) energy expenditure; cumulative data over the first 12 (C) and 24 (D) hours are also reported. Data are reported as mean ± SEM; no differences were detected between groups.

### Pain score

Pain was assessed using a rubric modified from one published by Adamson *et al*. and Clark *et al*. (Table 1).^14,17^ There were four categories: coat, eyes, coordination, and overall condition. Each category was allotted scores from 0-5, 0 being normal (no pain or distress), and 5 being the highest level of pain and distress. All animals were scored before the surgery, and scores were all zero (data not shown). Then, all animals were scored by a blinded scorer 6 h post-surgery, then once a day until 3 d post-surgery. The results showed that overall, Sham+C had the lowest pain scores. At 6 h post-surgery, Sx+C scored significantly higher than both Sham+C and Sx+C+SRB (Fig. 6). At 1 d post-surgery, both Sx+C and Sx+C+SRB scored significantly higher than Sham+C. These data suggest that neither carprofen-only nor carprofen and SRB analgesia regimen provided adequate pain control during the first day post-surgery.

**Figure 6.**
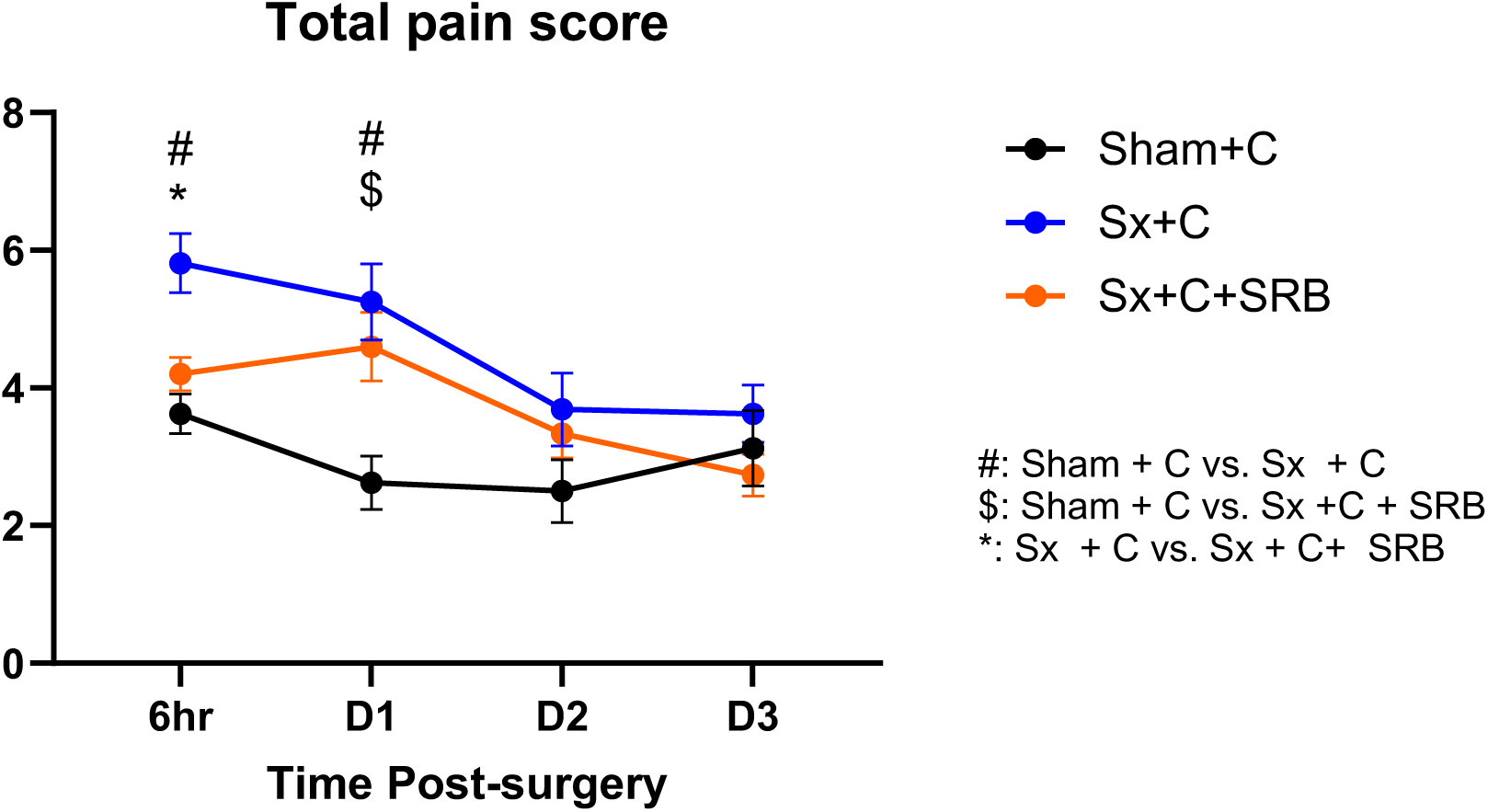
Pain scores. Total pain scores were calculated according to the rubric in Table 1. Data were recorded 6 h after surgery and for 1, 2, and 3 days after surgery (D1, D2, and D3, respectively). Data are reported as mean ± SEM. # *p* < 0.05 for Sham+C vs. Sx+C; $ *p* < 0.05 for Sham+C vs. Sx+C+SRB; * *p* < 0.05 for Sx+C vs. Sx+C+SRB.

**Table 1.** Post-surgical pain scores were determined and calculated according to the following rubric.

| Score | Coat | Eyes | Coordination | Overall condition |
| --- | --- | --- | --- | --- |
| 0 | Normal, well groomed, smooth sleek hair coat | Open, alert | Normal | Normal |
| 1 | Not well groomed | Squinted eyes | Walk awkward or slightly hunched; still runs or moves about | Rough appearance but acts fairly normal |
| 2 | Rough hair coat, dirty incision | Closed | Walk hunched, walking on eggshells, doesn't run, walks slowly | Slightly depressed, rough appearance or slightly agitated |
| 3 | Very rough hair coat, hair loss, dirty incision |  | Walks slowly with effort | Very rough or very agitated |
| 4 |  |  | Hunched, stumbles when walking |  |
| 5 |  |  | Hunched, not moving |  |
| **Total Pain Score = Coat + Eyes + Coordination + Overall condition**<br>**Minimum Total Pain Score = 0; Maximum Total Pain Score = 13** |  |  |  |  |

## Discussion

Post-operative analgesia is an essential component of veterinary care for laboratory animals. Beyond being the most humane approach, post-operative analgesia is important for experimental rigor in that it minimizes variability due to ranging levels of distress. Despite the necessity of analgesia, its comprehensive effects on the physiological and behavioral responses of animal models, including rodents, is not fully characterized. In rats and humans, opioid administration is often associated with GI disturbances.^9,11^ Therefore, we expected to observe decreased food consumption in the mice that received SRB. However, rather than decreased food consumption, SRB-treated mice consumed more food during the first 12 h post-surgery; food consumption was similar between all groups during the remainder of the 72 h of post-surgery observation. Contrary to our original hypothesis, the data suggest that SRB treatment does not significantly disturb the GI system of the mice in our study.

We identified with direct quantitative data that SRB caused transient hyperactivity in C57BL/6J mice during the initial 12 h after jugular vein and carotid artery catheterization surgery. This acute, SRB-mediated hyperactivity was consistent with previous reports that opioids have a stimulatory effect.^7,8^ For the first time, buprenorphine-induced hyperactivity was captured by a comprehensive metabolic caging system in our study. The hyperactivity was short-lived, mostly concentrated during the initial 12 h post-surgery. Investigators who conduct behavioral, metabolic, and other related studies should keep the buprenorphine-induced short-term hyperactivity in mind when designing experiments. To our surprise, hyperactivity in the Sx+C+SRB group did not provoke higher energy expenditure. Perhaps the hyperactivity was too transient and/or the magnitude was too low to manifest heightened energy expenditure.

Nevertheless, the lack of increased energy expenditure is important to document for metabolic studies, as it negates the need for additional nutritional support in mice receiving post-surgical analgesia via buprenorphine.

The pain score rubric we used was modified from a previously work by Clark *et al*. and Adamson *et al*.^14,17^ Similar to our observation, Adamson *et al*. also observed higher pain scores in buprenorphine-treated mice after mammary fat pad removal. In both cases, the increase in pain score observed in buprenorphine-treated mice was statistically significant, albeit mild. Pain assessment in mice is intrinsically difficult. In many published studies, including the mouse grimace scale and the ones that used the original pain scoring system of which we modified, recorded videos or photographs of the animals for pain assessment to avoid disturbing the animals are required. However, it was not feasible to record videos or photographs in the current study because the metabolic cages were seated inside an environmentally-controlled cabinet as part of the system’s requirement. Although Sx+C and Sx+C+SRB both scored significantly higher in pain score than Sham+C during the first day post-surgery, the activity and food consumption data showed that multimodal pain management of carprofen plus SRB was likely sufficient in post-surgical pain. Sx+C tended to have lower food consumption and activity level than the other two groups, albeit not always significant, therefore it is likely that carprofen alone was not providing sufficient pain control.

There are several limitations in this study. First, our experimental design did not include a Sham+SRB group, which could have served as another control to demonstrate if Sham+SRB mice also exhibited hyperactivity compared to Sham+C. A second limitation is the method of pain assessment. Due to the mice being in metabolic cages, only visual assessment could be performed. Although pain assessment is not the primary focus in this study, the pain scoring results prompted a broader and ongoing discussion of how to perform cage-side assessment of pain in mice. Buprenorphine-treated mice scoring higher on a similar pain score has been observed before^14^, therefore it is possible that the particular pain scoring system is not suitable for buprenorphine-treated mice. When assessing pain in a surgical model, using a non-surgical or sham surgery group as the baseline is a common practice. However, it may be unrealistic to expect an animal subjected to surgery with adequate pain control to behave and appear exactly the same as an animal that did not receive the same surgery. A previous study reported that mice that received carotid artery catheterization were less likely to integrate new nesting material into an existing nest compared to mice that received no surgery,^18^ even when buprenorphine was administered for pain control. A deeper review of how to select appropriate pain assessment method in mice is outside the scope of the current study, but our results do add to the knowledge that evaluating pain in mice is not simple, and investigators using research animals should always consult with a laboratory animal veterinarian regarding pain management.

The current study only evaluated C57BL/6J mice, therefore future studies in different strains or different sources of mice are needed to learn the effect of strain and source on buprenorphine-induced hyperactivity. Despite the limited scope of the current study, our quantitative documentation of the transient, SRB-induced hyperactivity in mice is important to consider when designing metabolic and behavioral studies as well as in post-operative monitoring. Importantly, based on our indirect calorimetry data used to quantify energy expenditure, this transient hyperactivity should not require additional nutritional supplementation to prevent unwanted metabolic disturbances. In conclusion, we recommend that researchers should consider the ways that analgesics can affect metabolic and behavioral parameters, and clearly report the analgesia regimens in publications.

## Acknowledgements

We thank the staff of the Comprehensive Metabolic Phenotype Core and the Center for Comparative Medicine for their contributions to this study.

## References

1. Heidbreder C, Fudala PJ, Greenwald MK. History of the discovery, development, and FDA-approval of buprenorphine medications for the treatment of opioid use disorder. Drug Alcohol Depend Rep. 2023;6:100133. doi:10.1016/j.dadr.2023.100133

2. FDA. Approval letter(s): Subutex (Buprenorphine HCL) Suboxone (Buprenorphine HCL & Naloxone HCL Dihydrate) Tablets. Published online October 8, 2002.

3. *U.S*. Government Principles for the Utilization and Care of Vertebrate Animals Used in Testing, Research, and Training. Vol 50.; 1985:20741–20880.

4. Guide for the Care and Use of Laboratory Animals. 8th ed.; 2011. doi:10.17226/12910

5. Foley PL. Current options for providing sustained analgesia to laboratory animals. Lab Anim. 2014;43(10):364–371. doi:10.1038/laban.590

6. *Recognition and* Alleviation of Pain in Laboratory Animals.; 2009. doi:10.17226/12526

7. Cowan A, Doxey JC, Harry EJ. The animal pharmacology of buprenorphine, an oripavine analgesic agent. Br J Pharmacol. 1977;60(4):547–554. doi:10.1111/j.1476-5381.1977.tb07533.x

8. Falcon E, Maier K, Robinson SA, Hill-Smith TE, Lucki I. Effects of buprenorphine on behavioral tests for antidepressant and anxiolytic drugs in mice. Psychopharmacology (Berl). 2015;232(5):907–915. doi:10.1007/s00213-014-3723-y

9. Clark JAJ, Myers PH, Goelz MF, Thigpen JE, Forsythe DB. Pica behavior associated with buprenorphine administration in the rat. Lab Anim Sci. 1997;47(3):300–303.

10. Gurtu S. Mu receptor-serotonin link in opioid induced hyperactivity in mice. Life Sci. 1990;46(21):1539–1544. doi:10.1016/0024-3205(90)90427-s

11. Botz-Zapp CA, Foster SL, Pulley DM, et al. Effects of the selective dopamine D(3) receptor antagonist PG01037 on morphine-induced hyperactivity and antinociception in mice. Behav Brain Res. 2021;415:113506. doi:10.1016/j.bbr.2021.113506

12. Camilleri M, Lembo A, Katzka DA. Opioids in Gastroenterology: Treating Adverse Effects and Creating Therapeutic Benefits. Clin Gastroenterol Hepatol Off Clin Pract J Am Gastroenterol Assoc. 2017;15(9):1338–1349. doi:10.1016/j.cgh.2017.05.014

13. Saenz M, Bloom-Saldana EA, Synold T, Ermel RW, Fueger PT, Finlay JB. Pharmacokinetics of Sustained-release and Extended-release Buprenorphine in Mice after Surgical Catheterization. J Am Assoc Lab Anim Sci JAALAS. 2022;61(5):468–474. doi:10.30802/AALAS-JAALAS-22-000025

14. Adamson TW, Kendall LV, Goss S, et al. Assessment of carprofen and buprenorphine on recovery of mice after surgical removal of the mammary fat pad. J Am Assoc Lab Anim Sci JAALAS. 2010;49(5):610–616.

15. Ayala JE, Bracy DP, Malabanan C, James FD, Ansari T, Fueger PT, McGuinness OP, Wasserman DH. Hyperinsulinemic-Euglycemic Clamps in Conscious, Unrestrained Mice. J Vis Exp. 2011;(57):3188. doi: 10.3791/3188.

16. Mina AI, LeClair RA, LeClair KB, Cohen DE, Lantier L, Banks AS. CalR: A Web-Based Analysis Tool for Indirect Calorimetry Experiments. Cell Metab. 2018;28(4):656–666.e1. doi:10.1016/j.cmet.2018.06.019

17. Clark MD, Krugner-Higby L, Smith LJ, Heath TD, Clark KL, Olson D. Evaluation of liposome-encapsulated oxymorphone hydrochloride in mice after splenectomy. Comp Med. 2004;54(5):558–563.

18. Gallo MS, Karas AZ, Pritchett-Corning K, Garner Guy Mulder JP, Gaskill BN. Tell-tale TINT: Does the Time to Incorporate into Nest Test Evaluate Postsurgical Pain or Welfare in Mice? J Am Assoc Lab Anim Sci JAALAS. 2020;59(1):37–45. doi:10.30802/AALAS-JAALAS-19-000044

